# Quantitative image analysis reveals no sexual dimorphism in the cellular dynamics of *Drosophila* heart tube formation

**DOI:** 10.64898/2026.04.23.720323

**Authors:** Rafael Perez-Vicente, Negar Balaghi, Rodrigo Fernandez-Gonzalez

**Affiliations:** Institute of Biomedical Engineering, University of Toronto, Toronto, ON, M5S 3G9, Canada; Translational Biology and Engineering Program, Ted Rogers Centre for Heart Research, University of Toronto, Toronto, ON, M5G 1M1, Canada; Division of Engineering Science, University of Toronto, Toronto, ON, M5S 1A4, Canada; Department of Cell and Systems Biology, University of Toronto, Toronto, ON, M5S 3G5, Canada; Developmental and Stem Cell Biology Program, The Hospital for Sick Children, Toronto, ON, M5G 1X8, Canada

## Abstract

Congenital heart defects affect females and males differently. Several congenital heart defects arise during the formation of the heart tube, suggesting that heart tube morphogenesis may differ between females and males. We investigated if the fruit fly *Drosophila melanogaster* displays sexual dimorphisms in the cellular mechanisms of heart tube formation. Quantitative microscopy revealed no differences between females and males in the migration of cardiac progenitors to form the heart tube. Our results suggest that *Drosophila* do not display sexual dimorphisms in early cardiac development, and support the omission of sex as an experimental variable when investigating *Drosophila* heart tube morphogenesis.

## Description

Congenital heart defects (CHDs) remain the leading birth defect (Zimmerman et al., 2020; Dotson et al., 2024). Anomalies in the early stages of heart development are associated with embryonic lethality and several CHDs (Gittenberger-de Groot et al., 2005; Bruneau, 2008; Ivanovitch et al., 2017). This includes outflow tract malformations, which account for 30% of CHDs (Sinha et al., 2015). A clear sexual dimorphism exists on the incidence of CHDs and outflow tract defects in particular (Mercuro et al., 2014; Deegan et al., 2021; Pugnaloni et al., 2023). Abnormal heart tube development is often a consequence of the defective migration of cardiac progenitors (Kelly et al., 2001; Sinha et al., 2015). Whether differences in the cellular dynamics of heart tube formation underlie the sexual dimorphisms in CHD has not been investigated.

The fruit fly *Drosophila melanogaster* is an excellent system to investigate the dynamics of cardiac morphogenesis. In both vertebrates and invertebrates heart development begins with the formation of a tube (Bodmer, 1995; Moorman & Christoffels, 2003; Souidi & Jagla, 2021). The heart tube forms as two groups of cardiac progenitors move toward each other from opposite sides of the embryo. In *Drosophila*, two contralateral rows of 52 cardiac progenitors (cardioblasts) each, migrate toward the dorsal midline, where they meet and form a tube (McFaul & Fernandez-Gonzalez, 2017). We showed that cardioblasts do not move monotonically forward (Balaghi et al., 2023). Instead, cardioblasts take cyclic forward and backward steps associated with waves of the molecular motor non-muscle myosin II and periodic cell shape changes. Periodic cardioblast steps are rectified into directional movement by a supracellular cable formed by the cytoskeletal protein filamentous actin (F-actin) at the trailing edge of cardioblasts. Potential differences in the cellular dynamics of heart tube formation between female and male *Drosophila* embryos have not been investigated.

To establish if there are differences in tissue dynamics between female and male *Drosophila* embryos during heart tube formation, we imaged heart tube closure in embryos expressing the cardiac-specific F-actin reporter hand-GFP:MoesinABD (Haack et al., 2014), and sxl-Pe:GFP (Thomson et al., 2004), which displays female-specific cytoplasmic GFP expression (Fig. 1A-A’’, 0 min). Visual inspection did not reveal any significant differences in dynamics of heart tube formation between females and males (Fig. 1A’’, B and Video S1). We used a recurrent neural network architecture, ReSCU-Net (Hawkins et al., 2025), to automatically trace the outline of the heart. We quantified luminal area changes over time (Fig. 1C) and fitted an exponential decay to the luminal area measurements to extract a heart tube closure rate constant (Fig. 1D). We found that the heart tube closed with a rate constant of 1.4±0.1 hr^-1^ (mean±standard deviation, s.d.) in female embryos, similar to the closure rate constant of 1.2±0.1 hr^-1^ in male embryos (Fig. 1D). To further determine if there were differences in the morphology of the heart tube as it formed, we quantified the circularity and maximum length of the luminal area along the anterior-posterior and medial-lateral axes of the embryo (Fig. 1E-F). Consistent with our measurements of heart tube closure dynamics, we found no differences in heart tube morphology between female and male embryos. Thus, our results suggest no sexual dimorphism exists in the tissue dynamics of *Drosophila* heart tube closure.

**Figure 1.**
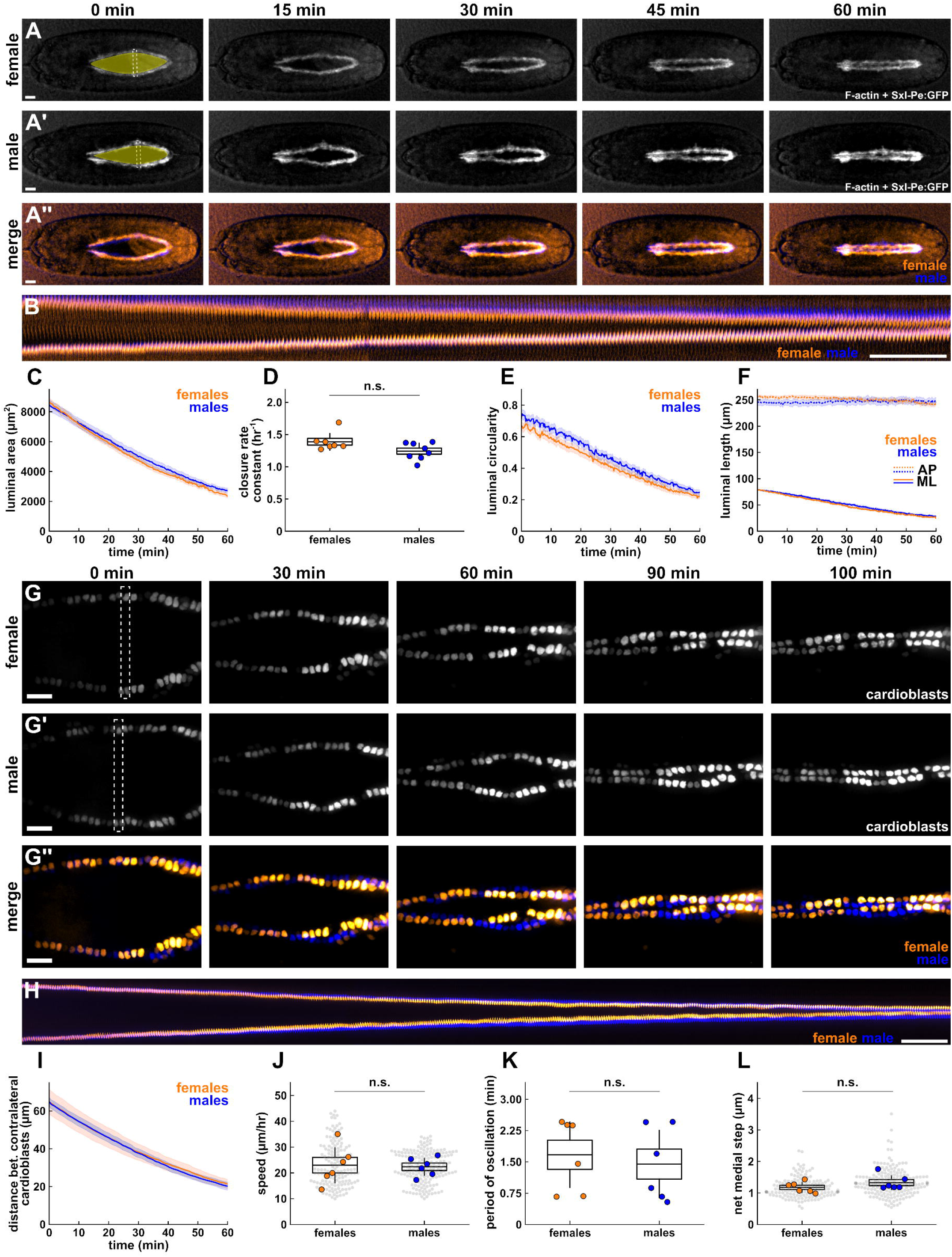
The tissue and cell dynamics of *Drosophila* heart tube formation show no sexual dimorphism. **(A, G)** Heart tube closure (A) and cardioblast migration (G) in female (A, G, orange in A’’, G’’) or male (A’, G’, blue in A’’, G’’) embryos expressing the heart-specific F-actin reporter hand-GFP:MoesinABD and the female-specific reporter sxl-Pe-GFP (A), or the cardioblast-specific nuclear marker Mid^E19^:GFP (G). The dominant orange hue outside the heart in (A’’) indicates sxl-Pe-GFP expression. Shading in (A-A’) indicates the luminal area. Dashed rectangles show the regions used to generate the kymographs in (B, H). **(B, H)** Kymographs of the dashed regions in (A) and (G), respectively. **(C-F)** Luminal area (C), heart tube closure rate constant (D), luminal circularity (E), and maximum anterior-posterior and medial-lateral luminal lengths (F) for females (orange, *n* = 7 embryos) or males (blue, *n* = 8). **(I-L)** Distance between contralateral pairs of cardioblasts (I), cardioblast speed (J), period of oscillation (K), and net medial step (L) in females (orange, 152 cardioblasts in *n* = 6 embryos), or males (blue, 184 cardioblasts in *n* = 6 embryos). (A, B, G, H) Anterior, left; medial, centre. Bars, 20 μm (A, G) or 5 min (B, H). (C, E-F, I) Error bars show the standard error of the mean. (D, J-K) Line indicates the mean, boxes show the standard error of the mean, bars indicate the standard deviation.

To further determine if there are differences in the cellular behaviours associated with heart tube formation in *Drosophila*, we measured cardioblast migration dynamics during heart tube development (Fig. 1G-H and Video S2). We imaged heart tube closure in embryos expressing a cardioblast-specific nuclear marker, Mid^E19^:GFP (Jin et al., 2013). We previously showed that nuclear dynamics match cellular dynamics in *Drosophila* cardioblasts (Balaghi et al., 2023). After imaging, embryos were allowed to develop into larvae and adults, and were sexed based on gonad or body morphology (Hartenstein, 1993). We automatically outlined cell nuclei from confocal microscopy movies using a U-Net, a convolutional neural network architecture (Falk et al., 2019; Balaghi et al., 2023). We found that cardioblasts migrated towards the dorsal midline at a similar speed in both females and males (23.0±6.7 μm/hr *vs*. 22.4±3.3 μm/hr, respectively, Fig. 1I-J). To further establish if the mechanisms of cell movement varied between females and males, we quantified the oscillatory dynamics of cardioblast nuclei. Our results showed that the period of cardioblast oscillation was not significantly different between females and males (1.7±0.8 min *vs*. 1.4±0.8 min, respectively, Fig. 1K). We could not detect any differences in the amplitude of the medial (1.9±0.1 μm for females *vs*. 2.0±0.2 μm for males) or lateral (0.8±0.1 μm for females *vs*. 0.7±0.1 μm for males) steps, with comparable net medial steps of 1.2±0.2 μm in females and 1.3±0.2 μm in males (Fig. 1L). Thus, our results indicate that there are no sexual dimorphisms in the dynamics of cardioblast migration during *Drosophila* heart tube formation.

Altogether, our results suggest that the early stages of heart development in *Drosophila* do not display sexual dimorphisms, at least not at the level of cellular or tissue dynamics. The oscillatory movement of cardioblasts and the coordination of cardioblast migration is driven by periodic myosin waves and a supracellular F-actin cable (Balaghi et al., 2023). Thus, our results suggest that the cytoskeletal structures associated with heart tube formation will be similar between females and males. This is consistent with evidence from other systems in which actomyosin-based cytoskeletal networks drive collective cell movement. For example, the wound healing response in monolayer epithelia of both female and male origin is associated with a supracellular actomyosin cable that guides and coordinates cell movements (Bement et al., 1993; Marpeaux et al., 2026). Our data support the omission of sex as a biological variable in studies of cellular dynamics during *Drosophila* heart tube formation. However, a limitation of our assay is that we did not quantify cell or tissue dynamics in naturally-occurring abnormal hearts (e.g. missing cardioblasts). Further studies measuring the relative incidence of defective heart tubes in females and males, at embryonic, larval, and adult stages, will reveal the extent to which *Drosophila* can be used to model sexual dimorphisms in CHDs.

## Methods

### Fly stocks

We used Flybase (release FB2025_03) to find information on stocks and gene expression (Öztürk-Çolak et al., 2024). *D. melanogaster* strains (see Resources and Reagents) were kept at room temperature on fly food provided by a central kitchen operated by H. Lipshitz. Embryos were collected overnight on apple juice agar plates in plastic cages at room temperature (∼23°C). Stage 14-15 embryos (collected 12-14 hours after egg laying) were used.

We used *mid*^*E19*^*:GFP* (Jin et al., 2013) to image cardioblast nuclei and *hand-GFP:moesinABD* (Haack et al., 2014) to visualize F-actin in the heart. *sxl-Pe-GFP* (Thomson et al., 2004) (Bloomington Drosophila Stock Center #32565) was used to identify female embryos based on the expression of cytoplasmic GFP driven by the female-specific *sxl-pe* promoter.

### Timelapse imaging

Embryos were dechorionated in 50% bleach in water for 2 minutes, rinsed and glued dorsal side down on a glass coverslip using heptane (Caledon Laboratory Chemicals) glue. Embryos were covered in a 1:1 mix of halocarbon oil 27 and 700 (Sigma-Aldrich) for live imaging (Scepanovic et al., 2021). We imaged embryos on a Revolution XD spinning disk confocal microscope (Andor) equipped with an iXon Ultra 897 camera, a 10x air lens (NA 0.40, Olympus) and a 60x oil-immersion lens (NA 1.35, Olympus). We acquired sixteen-bit Z-stacks every 15 seconds at 0.5 or 0.75 µm steps (21 to 37 slices per stack) using Metamorph Software (Molecular Devices). Maximum intensity projections of the stacks were generated with Fiji (Schindelin et al., 2012) and used for analysis.

### Sex classification

Embryonic sex was determined either by using sxl-Pe-GFP expression to identify female embryos prior to imaging, or by allowing embryos to develop after imaging and using gonad morphology in the larva and both gonad and body morphology in the adult to assign a sex to the corresponding movie (Hartenstein, 1993). Individual embryos were transferred to yeasted apple agar plates after imaging and allowed to develop at room temperature until third instar larva (5-6 days) or adult (12-14 days) stages.

### Image quantification

Automated object segmentation, editing and analysis were performed using our open-source image analysis platform, PyJAMAS (Fernandez-Gonzalez et al., 2022). For object segmentation, the LiveWire algorithm (Fernandez-Gonzalez & Zallen, 2013) was used to semi-automatically annotate heart lumens and cardioblast nuclei. We used semi-automated annotations to train a ReSCU-Net and a U-Net for luminal and nuclear segmentation, respectively (Falk et al., 2019; Hawkins et al., 2025). The heart tube closure rate constant, *k*, was defined by fitting the luminal area over time with a decaying exponential:

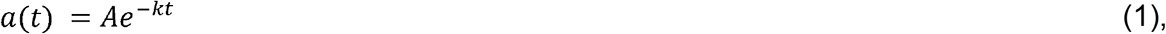

where *a*(*t*) is the luminal area at time *t*, and *A* is the maximum luminal area. We quantified the maximum AP and ML luminal lengths by rotating the luminal annotations such that the longest axis was perpendicular to the horizontal axis, and measuring the difference between the maximum and minimum x-coordinates (ML length) or y-coordinates (AP length). Luminal circularity at time *t, c*(*t*), was calculated as:

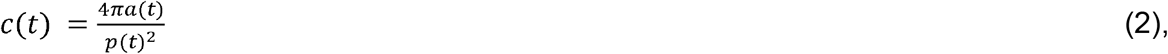

where *p*(*t*) represents the luminal perimeter at time *t*. Cardioblast positions were determined using the centroid of the cardioblast nucleus (Balaghi et al., 2023). The speed of cardioblast migration was the slope of a line fitted to the distance travelled by individual nuclei over time. The period of oscillation was measured as the time lag of the first peak of the autocorrelation of each cardioblast velocity curve.

### Statistical analysis

Embryonic means were compared using non-parametric Mann-Whitney tests. For time series, error bars indicate the standard error of the mean (s.e.m.). In box plots, error bars show the s.d., boxes indicate the s.e.m., and a line inside the box denotes the mean.

## Acknowledgments

We are grateful to Ana Maria do Carmo, Alexandra Korolov, Willow Peterson and Ji Hong Sayo for critical reading of this manuscript. RPV was partially supported by the Undergraduate Summer Research Program of the Ted Rogers Centre for Heart Research Translational Biology and Engineering Program. This work was funded by grants to RFG from the National Sciences and Engineering Research Council of Canada (RGPIN-2025-04527), the Canada Foundation for Innovation (30279 and 43988) and Ted Rogers Centre for Heart Research Translational Biology and Engineering Program. RFG is the Tier II Canada Research Chair in Quantitative Cell Biology and Morphogenesis.

## Resources and reagents

| Reagent or resource | Source | Identifier |
| --- | --- | --- |
| **Experimental models: organisms/strains** |  |  |
| *D. melanogaster: mid<sup>E19</sup>:GFP* | (Jin et al., 2013) | N/A |
| *D. melanogaster: hand-GFP:moesinABD* | (Haack et al., 2014) | N/A |
| *D. melanogaster: sxl-Pe-GFP* | (Thomson et al., 2004), Bloomington Drosophila Stock Center | 32565 |
| **Chemicals** |  |  |
| Heptane | Caledon Laboratory Chemicals | Cat #: 5400-1-10 |
| Halocarbon oil 27 | Sigma Aldrich | Cat #: 118773 |
| Halocarbon oil 100 | Sigma Aldrich | Cat #: 118898 |
| **Software** |  |  |
| PyJAMAS | (Fernandez-Gonzalez et al., 2022) | pyjamas.readthedocs.io |
| Fiji | (Schindelin et al., 2012) | imagej.net |
| MetaMorph | Molecular Devices | RRID:SCR,_002368 |

## Video legends

**Video S1. Tissue dynamics during *Drosophila* heart tube formation are similar between female and male embryos**. Heart tube closure in a female (left, orange in merge) and a male (centre, blue in merge) embryo expressing the heart-specific F-actin reporter hand-GFP:MoesinABD and the female-specific reporter sxl-Pe-GFP. Images were acquired every 15 s. Anterior left; medial, centre.

**Video S2. Cardioblast dynamics during *Drosophila* heart tube formation are similar between female and male embryos**. Cardioblast migration in a female (left, orange in merge) and a male (centre, blue in merge) embryo expressing the cardioblast-specific nuclear marker Mid^E19^:GFP. Images were acquired every 15 s. Anterior left; medial, centre.

